# Melanoma-derived DNA polymerase theta variants exhibit altered DNA polymerase activity

**DOI:** 10.1101/2023.11.14.566933

**Authors:** Corey Thomas, Lisbeth Avalos-Irving, Jorge Victorino, Sydney Green, Morgan Andrews, Naisha Rodrigues, Sarah Ebirim, Ayden Mudd, Jamie B. Towle-Weicksel

**Affiliations:** Department of Physical Sciences, Rhode Island College, Providence, RI 02908

**Keywords:** <u>Keywords</u>: DNA polymerase, melanoma, mutagenesis mechanism, DNA repair, enzyme kinetics

## Abstract

DNA Polymerase θ (Pol θ or POLQ) is primarily involved in repairing double-stranded breaks in DNA through the alternative pathway known as microhomology-mediated end joining (MMEJ) or theta-mediated end joining (TMEJ). Unlike other DNA repair polymerases, Pol θ is thought to be highly error prone, yet critical for cell survival. We have identified several mutations in the POLQ gene from human melanoma tumors. Through biochemical analysis, we have demonstrated that all three cancer-associated variants experienced altered DNA polymerase activity including a propensity for incorrect nucleotide selection and reduced polymerization rates compared to WT Pol θ. Moreover, the variants are 30 fold less efficient at incorporating a nucleotide during repair and up to 70 fold less accurate at selecting the correct nucleotide opposite a templating base. Taken together, this suggests that aberrant Pol θ has reduced DNA repair capabilities and may also contribute to increased mutagenesis. While this may be beneficial to normal cell survival, the variants were identified in established tumors suggesting that cancer cells may use this promiscuous polymerase to its advantage to promote metastasis and drug resistance.

## Introduction

DNA is constantly damaged by endogenous and exogenous factors including free radicals, chemical agents, and ionizing and UV radiation. Endogenous damage alone is estimated to occur at a rate of at least 20,000 lesions per cell per day^1^. These lesions result in a variety of issues including the formation of abasic sites, thymine dimers, single strand breaks, and/or double strand breaks, which if left unrepaired, can lead to genomic instability, cancer, and/or cell death. Due to this high level of DNA damage, the cell employs DNA repair pathways including homologous recombination and non-homologous end joining to repair double strand breaks, and to maintain the stability of the genome. DNA polymerase theta, Pol θ, is the major DNA polymerase in the alternative double-stranded DNA repair pathway known as microhomology-mediated end joining (MMEJ) or theta-mediated end joining (TMEJ) ^2–4^. Unlike the more robust and precise repair of double strand breaks in the homologous recombination repair pathway which employs nearby sister chromatid as a template during the S and G2 cell cycle to ensure accuracy ^5,6^, TMEJ utilizes internal microhomologies of 2-6 base pairs within the DNA it is repairing as a template ^3,7^. Despite Pol θ being naturally error-prone repair enzyme^8,9^, it is hypothesized that its TMEJ pathway acts as an auxiliary repair method especially when other repair pathways are compromised ^10–13^. However, upregulation of *POLQ* has been demonstrated to be particularly harmful to patients especially those with lung and breast cancers ^14,15^ whereas knock-out studies of *POLQ* suggest sensitivity ionizing radiation and an increased likelihood of developing skin cancer ^4,9,16–19^. Furthermore, it is possible that Pol θ allows for cell survival despite losing genomic integrity to a certain extent ^20^. Taken together, this suggests that a cancer cell could take advantage of this low-fidelity DNA repair polymerase to promote drug resistance and metastasis^21^. Thus, we wanted to explore the mechanistic function of Pol θ during DNA repair.

Pol θ is a large A-family DNA polymerase (290 kDa) enzyme that contains a N-terminus helicase domain (residues 1-891) and a C-terminus polymerase domain (residues 1819-2590) tethered together by an unstructured central domain (residues 892-1818) ^17,22–27^. The C-terminus can be isolated and characterized as a fully functional DNA polymerase where it contains classic DNA polymerase subdomains including the DNA binding thumb (residues 2093- 2217), nucleotide binding fingers (residues 2333-2474), catalytic active site palm (residues 2218-2590), and an exonuclease-like domain (residues 1819-2090)^27,28^. While many studies have looked at the importance of overexpression or deficient *POLQ* in relation to genomic stability, little is known about the biochemical mechanism on a molecular level by which Pol θ selects and incorporates specific nucleotides during DNA. Sporadic and hereditary mutations have been found in all five DNA polymerase families expressed in a variety of tumors ^29–33^ .

Many of these cancer-associated variants have been characterized *in vitro* and suggest that aberrant DNA polymerases often experience slower polymerization rates, increased mutagenesis, and/or poor repair abilities past lesions which lead to chromosomal aberrations and genomic instability^33–40^ . Here we have identified several *POLQ* variants from melanoma patients from Tissue Resource Core of the Yale SPORE in Skin Cancer. Focusing specifically on the C-terminal DNA polymerase domain, we selected variants from each of the three subdomains (thumb, fingers, palm) that were predicted to be detrimental to the enzymes structure and function through SIFT and PolyPhen algorithms (Table 1, Figure 1)^41,42^. Mutations were introduced into the isolated C-terminal construct *PolQM1*^28^ for an in-depth analysis of deoxynucleotide affinity (Kd(dNTP)) and rates of DNA extension (*k_pol_*) to ultimately relate decreased fidelity to an increased state of genomic instability compared to wild-type Pol θ. All three variants (T2161I, E2406K, and L2538K) demonstrate a decreased fidelity and our study provides structural insight into key residues needed for DNA binding, nucleotide selection, and polymerization for accurate DNA repair.

**Figure 1.**
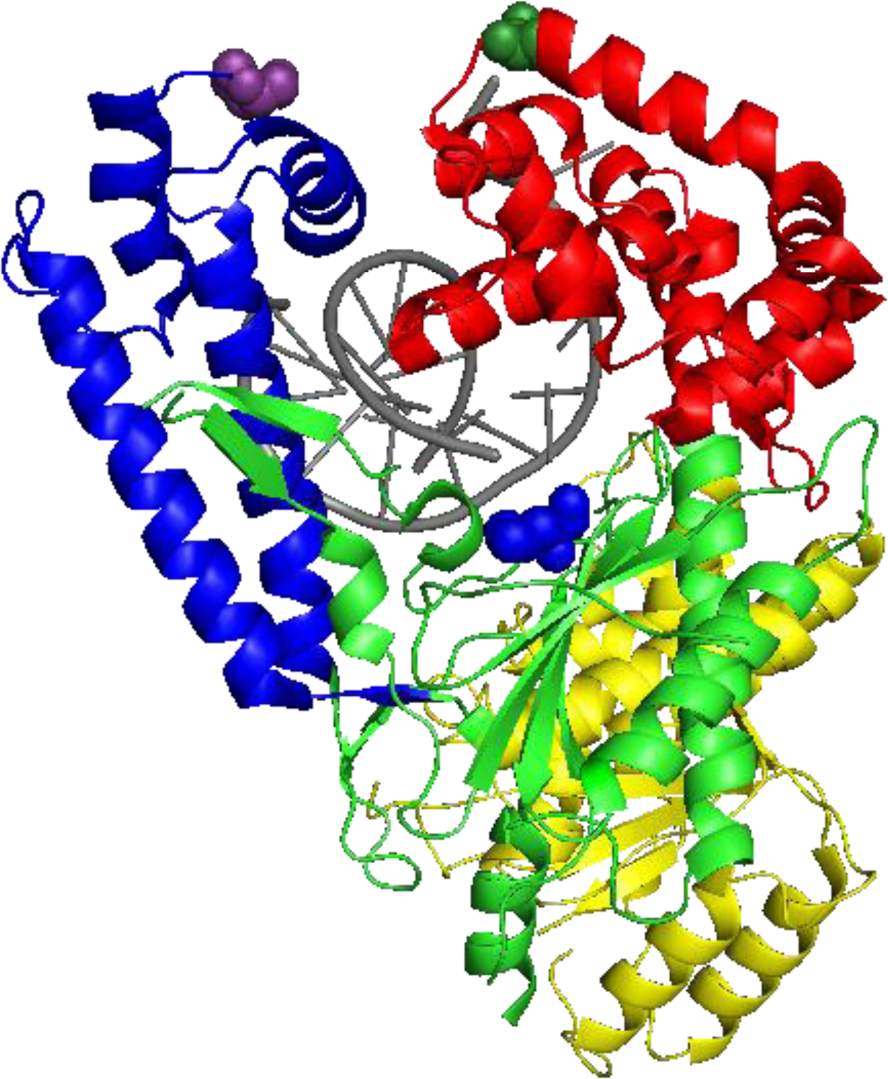
Missense amino acid substitutions found in the polymerase domain of DNA Pol θ. The truncated c-terminal polymerase domain of Pol θ has four subdomains: thumb (blue), fingers (red), palm (green), and exonuclease domain (yellow). The missense amino acid changes from melanoma patients are represented by spheres with T2161I in purple, E2406K in dark green, and L2538R in blue. The DNA substrate is indicated in gray (protein data bank code 4X0Q^26^.

**Table 1.** Missense amino acid substitutions in DNA polymerase theta from melanoma tumors with algorithm predictions. ^54^.

| **Variant** | **Pol θ** | **Melanoma** | **Stage** | **SIFT** | **PolyPhen-2** |
| --- | --- | --- | --- | --- | --- |
|  | **Subdomain** | **Type** |  |  |  |
| T2161I | Thumb | Sun-exposed | II | Deleterious | Probably Damaging |
| E2406K | Fingers | Ocular | IV | Deleterious | Probably Damaging |
| L2538R | Palm | Sun-exposed | IV | Deleterious | Probably Damaging |

## Results

### Twelve percent of Melanoma tumors contained mutations in POLQ

Of the 250 melanoma samples obtained from the Yale SPORE in Skin Cancer, 29 patients had at least one mutation in the POLQ gene resulting in 11.6% occurrence. From these 29 patients, 9 had missense mutations in the c-terminus Polymerase Domain defined between 1792-2590^28^. Choosing a cancer-associated variant from each of the three major DNA polymerase subdomains; thumb, fingers, and palm, we looked to identify key residues that are involved in nucleotide incorporation and fidelity (Figure 1, Table 1).

The mutations were introduced individually into the human pSUMO3 c-terminal POLQM1 via site-directed mutagenesis^28^. Constructs were expressed and purified from *E.coli* with an average final concentration of 3μM. To ensure individual point mutations did not affect the overall structure, circular dichroism spectroscopy was performed on WT and variants at 20°C. The spectra of each variant were similar to that of WT Pol θ suggesting that all variants had similar secondary structure characteristics (Figure 2).

**Figure 2.**
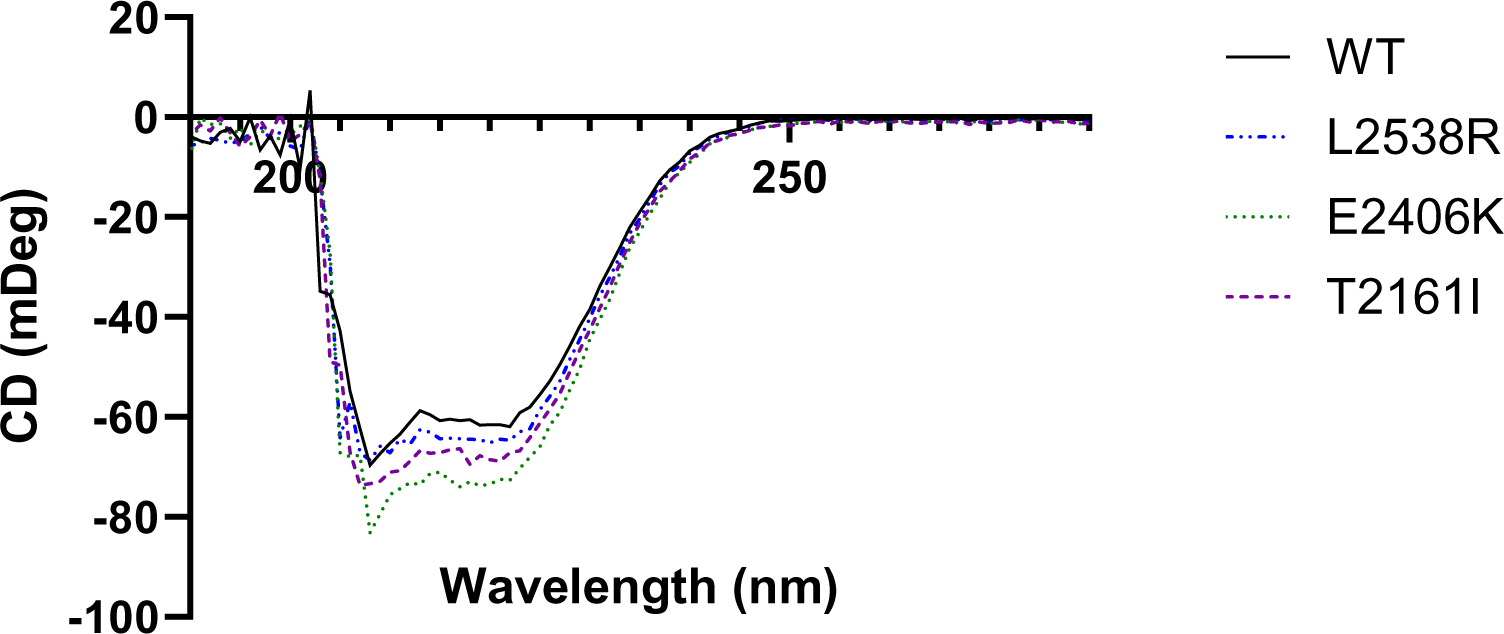
Secondary structure of cancer-associated variants similar to WT Pol θ. Circular dichroism spectra of 3μM WT (black, solid line) and variants L2538R (blue, dot and dashed line), E2406K (green, small dashed line) and T2161I (purple, large dashed line) in 10mM K_2_HPO_4_ scanned from 190 to 280nm at 20°C.

### Pol θ variants experience biphasic burst kinetics

To assess the overall DNA polymerase activity, all variants and WT Pol θ were assayed under pre-steady state burst kinetics (Figure 3 and Table 2). Variant T2161I had amplitude and rate limiting *k_ss_* kinetics compared to WT but was twice as fast with an observed rate (*k_obs_) of 142 s^-^*^1^. Variants E2406K and L2538R displayed a greater amplitude but had a 2-fold slower observed rate, around 28-38 s^-1^. For all variants, the product extension fit to a biphasic burst equation (equation 2) with the rate limiting step being product release, as observed in WT Pol θ.

**Figure 3.**
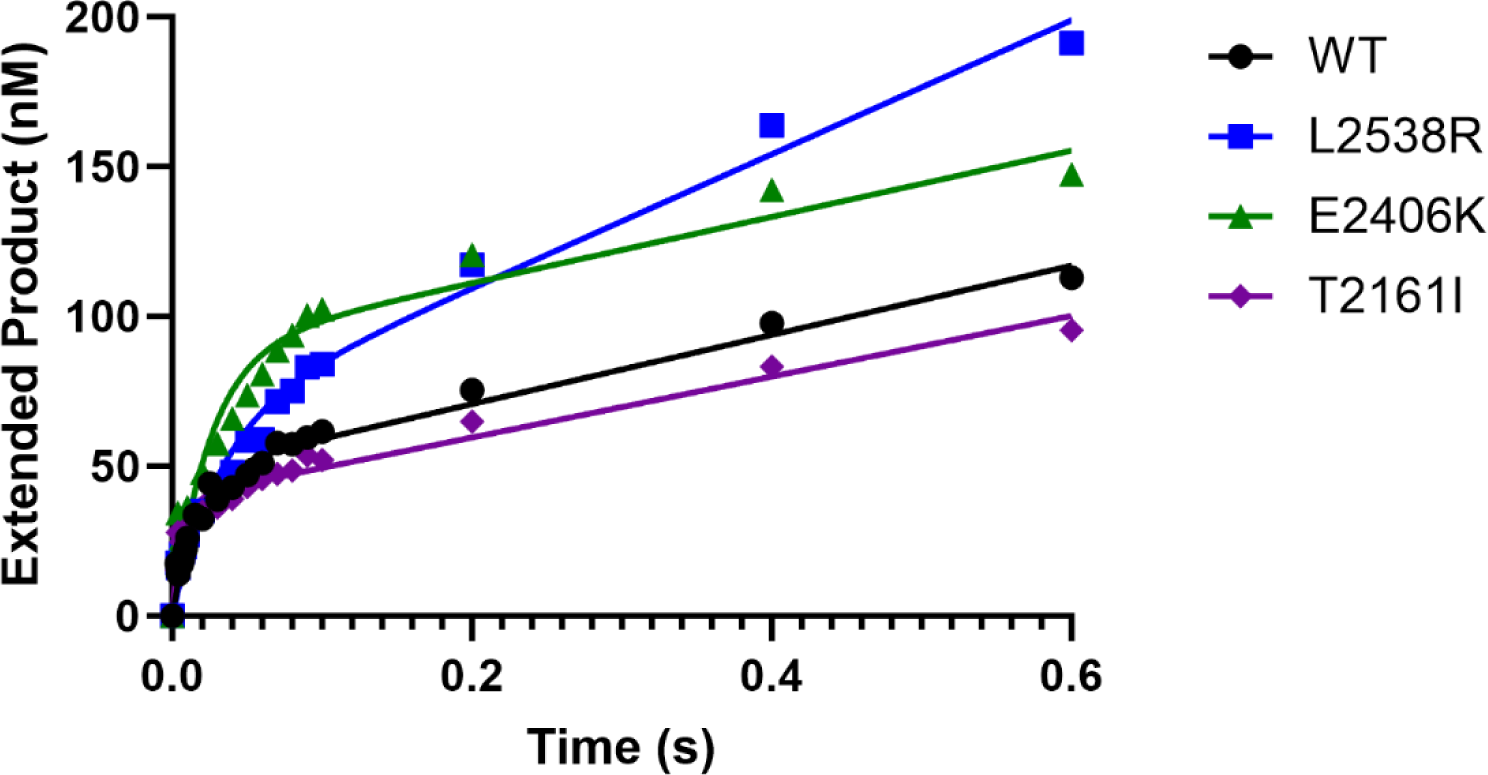
Wild-type and cancer-associated variants experience fast biphasic burst kinetics. Pre-steady state burst kinetics were performed by pre-incubating 100nM duplex DNA with 300nM of WT (circles), T2161I (diamond), E2406K (triangles), or L2538R (squares) and reacting with 100μM dCTP (correct) from 0.0037 to 0.6s. Data was graphed as extended product versus time and fit to Equation 1.

**Table 2.** Observed polymerization rates of Pol θ and cancer-associated variants.

| Pol $\theta$ | $k_{obs}$ ($s^{-1}$) | $k_{ss}$ ($s^{-1}$) | $E_{app}$ (nM) | Fold change<sup>a</sup> |
| --- | --- | --- | --- | --- |
| WT | 60.00 ± 6.1 | 2.336 ± 0.2 | 51 ± 1.8 | 1 |
| T2161I | 142.00 ± 22.5 | 2.538 ± 0.3 | 41 ± 1.8 | 0.423 |
| E2406K | 38.12 ± 5.2 | 1.201 ± 0.2 | 95 ± 4.8 | 1.6 |
| L2538R | 27.96 ± 5.3 | 3.107 ± 0.2 | 80 ± 7.0 | 2.1 |

<sup>a</sup> $k_{obs}$ WT/ $k_{obs}$ variant

### Cancer-associated variants experience a different kinetic pathway compared to WT

In order to investigate the difference between the observed DNA polymerization rates between WT and variants, single turnover kinetics were carried out with excess protein (4:1, protein to DNA) to determine the polymerization rate (*k_pol_*) and binding affinity (K_d(dNTP)_) for each nucleotide opposite a DNA template G. Each variant and WT protein preparation was assayed for percent activity through an active site titration. The enzymes were assayed at varying DNA concentrations to determine the amplitude. This was plotted and mathematically fit to a quadratic equation. Each protein preparation was approximately 62% active and experimental protein concentrations were adjusted to equal active protein levels (Figure 4). Rates and affinity were verified empirically through enzyme titration to ensure maximum product formation (data not shown).

**Figure 4.**
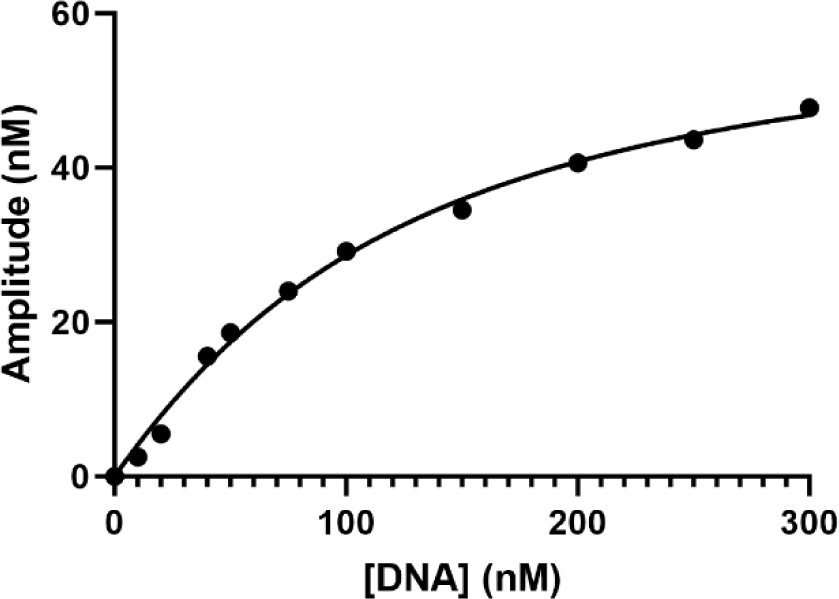
Representative Active site titration demonstrates 62% active protein. Duplex DNA was titrated from 0 to 300nM against 100nM Pol θ and 100μM dCTP 0.003 to 0.6s. Products were separated and visualized similar to the pre-steady state burst kinetic experiments and a E_app_ was calculated for each DNA concentration. Amplitude versus DNA was graphed to determine K_D(DNA)_ at 84.1 ± 9.5nM and Active sites at 62.4 ± 2.7%.

These experiments revealed polymerization rates similar to the pre-steady state burst assay. Both variants E2406K and L2538R experienced reduced *k_pol_* rates compared to WT with 89 s^-1^ and 41 s^-1^ respectively. Interestingly, under these conditions both WT and T2161I demonstrated a rapid polymerization rate of around 167s^-1^

For WT we observed a correct nucleotide affinity of 5μM whereas L2538R experienced a 3-fold less affinity for correct, E2406K an 18-fold decrease, and T2161I a 38-fold decrease.

Conversely, all variants preferred to incorporate the incorrect nucleotides dATP and dTTP, observed by the lower determined K_d_ value when compared to WT. Taking into account both *k_pol_* and K_d_, indicators of overall polymerase efficiency and fidelity, all variants experienced a reduction in these parameters compared to WT.

## Discussion

DNA polymerase’s biochemical kinetics can provide key insight into DNA damage repair. The fidelity, or ability of the DNA polymerase to select the correct nucleotide during nucleotide incorporation is an important step in DNA repair and preventing genomic instability and cancer. By determining the *k_pol_ and* K_d(dNTP)_ a more cohesive mechanism of polymerase activity can be observed, providing insight into how aberrant enzymes may alter this activity ^43^. DNA Polymerase θ has been shown to be a low fidelity enzyme that is critical for alternative double strand break repair especially in BRCA-depleted cells ^18,26,44^. What is unclear about the role of Pol θ in DNA repair is whether the polymerase stabilizes the genome through repair or if it detrimental to the cell to allow for increase mutagenesis. Perhaps reinforcing this apparent contradiction, prior studies have explored the role of POLQ in genomic stability in either loss of function or over-expression analyses; which have notable detrimental secondary effects^10,13^.

Additionally, prior to this study, catalytic mutations, without direct connection to disease, were examined in the both the N-terminal Helicase and C-terminal Polymerase domains for functional studies ^4,26,45–48^. Here we report for the first time, three Pol θ missense mutations reported in melanoma patients that biochemically display altered protein function, suggesting a potential role in genomic instability. These three mutations were strategically selected to explore the subdomains of the C-terminal polymerase domain of Pol θ in order to explore the functional role of each subdomain from a disease context. We generated these mutations *in vitro* and utilized classical primer extension assays to analyze nucleotide selection and incorporation compared to WT to gain insight into the variants overall DNA repair capabilities.

### Wild-type

Previous studies have explored pre-steady state kinetics of truncated Pol θ ^26,28^ and our studies observed similar rates. A common nucleotide incorporation mechanism for DNA polymerases can also be applied to Pol θ based on our biochemical data. Pol θ experiences a *k_obs_* around 60 ± 6 s^-1^ (steps including nucleotide binding, polymerization, and pyrophosphate release) that proceeds the slower rate limiting *k_ss_* rate (Figure 3, Table 2). Most kinetic studies of Pol θ have been performed under steady-state conditions^18^ which can underestimate the more transient nucleotide selection step that occurs just prior to polymerization^43^. To better capture factors such as *k_pol_* and K_d(dNTP)_, which directly influence fidelity and efficiency, we did our analysis using single-turnover conditions. Through this approach, we observe *k_pol_* of WT to be 167 s^-1^ with a K_d(dNTP)_ of around 5μM indicating tight binding to correct nucleotide (Figure 5, Table 3). As expected, incorrect nucleotide affinity was slightly higher, consistent with the steady-state assumption that Pol θ is a low fidelity enzyme. Particularly, WT incorporated dGTP opposite template G through very tight nucleotide binding. The explanation for this may be due to template slippage which has been seen in other DNA polymerases ^26,49^ due to an adjacent C in the DNA template, but also due to its affinity for purines as previously reported^7^. Overall, Pol θ experiences a faster polymerization rate than some higher fidelity DNA polymerases including Pol β, γ and ε, but tighter binding to correct nucleotide ^50–53^.

**Figure 5.**
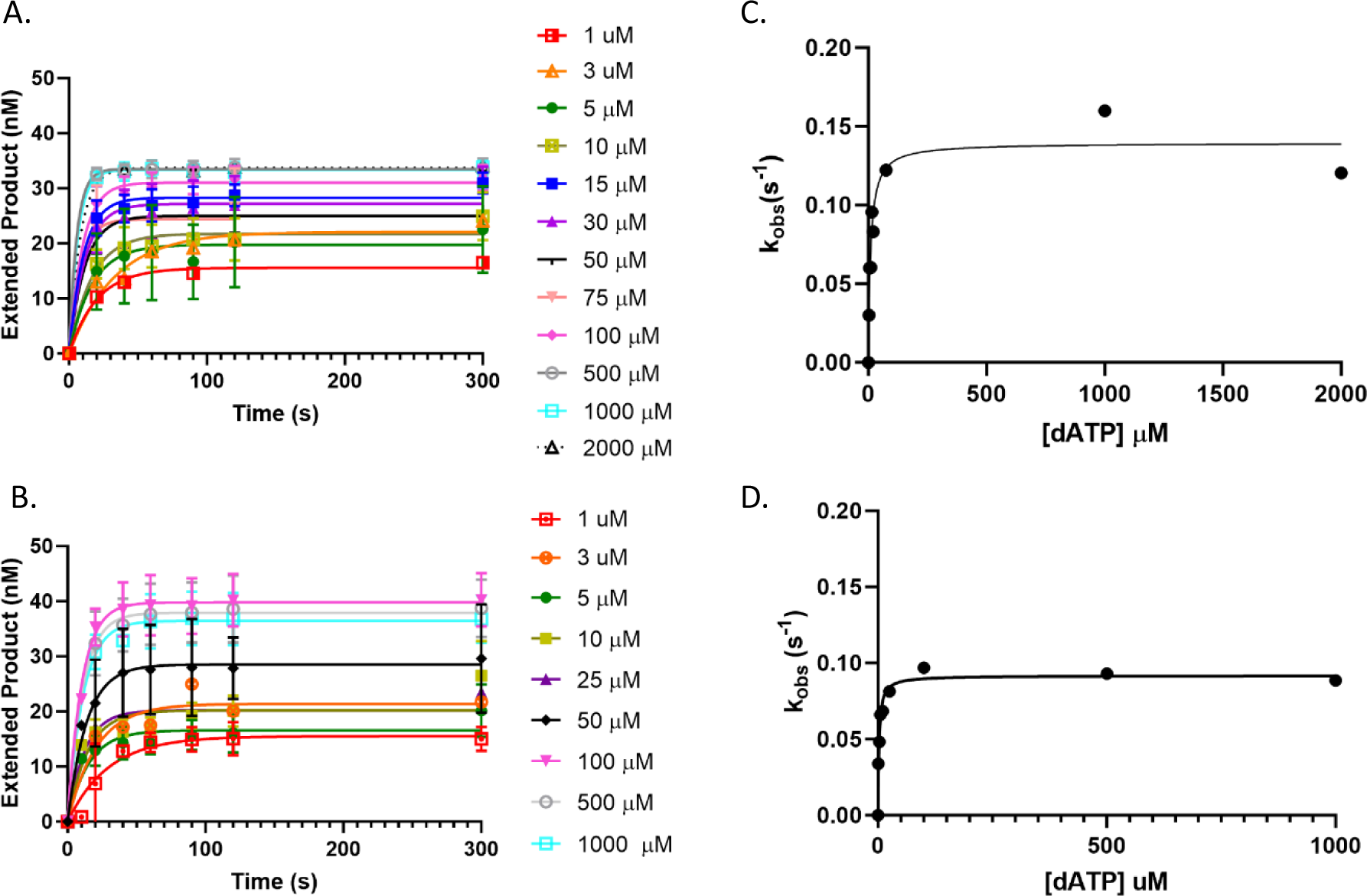
Cancer-associated variants bind to incorrect nucleotide with greater affinity than WT. A representative plot of single turnover experiments with incorrect dATP opposite template G. Increasing concentrations of dATP were titrated against 50nM DNA substrate and 200nM Pol θ (A) or L2538R (B). Extended product was graphed versus time to determine *k_obs_.* The observed rate was plotted against concentration of dATP (C and D) and data fit to a hyperbolic equation 4 to determine k_pol_ and K_d(dNTP)_.

**Table 3.** Single Turnover kinetics for WT Pol θ and Cancer-associated variants on duplex DNA with template G.

| Pol $\theta$ | Sequence | $k_{pol} (s^{-1})$ | $K_d (\mu M)$ | $D k_{pol}^a$ | $D K_d^b$ | Efficiency<sup>c</sup> | ΔEfficiency<sup>d</sup> | F<sup>e</sup> | $\Delta$ fidelity<sup>f</sup> |
| --- | --- | --- | --- | --- | --- | --- | --- | --- | --- |
| WT | G:dC | 167 ± 4.88 | 5.20 ± 0.68 | | | 32.14 | | | |
| WT | G:dG | 0.114 ± 0.004 | 1.60 ± 0.35 | 1466 | 0.31 | 0.07 | | 452 | |
| WT | G:dA | 0.139 ± 0.009 | 9.71 ± 2.4 | 1198 | 1.87 | 0.01 | | 2240 | |
| WT | G:dT | 0.150 ± 0.010 | 15.28 ± 3.7 | 1115 | 2.94 | 0.010 | | 3279 | |
| T2161I | G:dC | 166.5 ± 12.67 | 199.1 ± 41.7 | | | 0.84 | 38 | | |
| T2161I | G:dG | 0.223 ± 0.014 | 44.29 ± 9.8 | 746 | 0.22 | 5.04E-03 | 14 | 167 | 3 |
| T2161I | G:dA | 0.133 ± 0.008 | 15.18 ± 3.9 | 1255 | 0.08 | 8.74E-03 | 2 | 97 | 23 |
| T2161I | G:dT | 0.111 ± 0.005 | 5.50 ± 1.4 | 1507 | 0.03 | 2.01E-02 | 0.5 | 43 | 77 |
| E2406K | G:dC | 89.04 ± 5.44 | 93.08 ± 19.8 | | | 0.96 | 34 | | |
| E2406K | G:dG | 0.132 ± 0.006 | 17.38 ± 3.5 | 677 | 0.19 | 0.01 | 9 | 127 | 4 |
| E2406K | G:dA | 0.160 ± 0.009 | 33.83 ± 6.4 | 557 | 0.36 | 0.005 | 3 | 203 | 11 |
| E2406K | G:dT | 0.120 ± 0.007 | 10.17 ± 2.5 | 744 | 0.11 | 0.01 | 1 | 82 | 40 |
| L2538R | G:dC | 41.16 ± 1.07 | 15.70 ± 1.4 | | | 2.62 | 12 | | |
| L2538R | G:dG | 0.117 ± 0.004 | 3.50 ± 0.71 | 352 | 0.22 | 0.03 | 2 | 80 | 6 |
| L2538R | G:dA | 0.092 ± 0.003 | 2.29 ± 0.36 | 449 | 0.15 | 0.040 | 0.4 | 66 | 34 |
| L2538R | G:dT | 0.125 ± 0.008 | 12.08 ± 3.0 | 329 | 0.77 | 0.01 | 1 | 254 | 13 |

<sup>a</sup>(c/i) <sup>b</sup>(i/c) <sup>c</sup> $k_{pol}/K_d$ ( $\mu M^{-1}s^{-1}$ ) <sup>d</sup>Efficiency<sub>WT</sub>/Efficiency<sub>variant</sub> <sup>e</sup>F=(Efficiency<sub>correct</sub>+Efficiency<sub>incorrect</sub>)/Efficiency<sub>incorrect</sub> <sup>f</sup>Fidelity<sub>WT</sub>/Fidelity<sub>variant</sub>

### T2161I Thumb Domain mutation

The T2161I variant was originally discovered in a tumor from patient with stage 2 melanoma (Figure1, Table 1). Structure and function prediction algorithms suggested that this mutation, as well as the other two listed below, could be potentially deleterious and prompted further investigation into its biochemical function. Like all of the cancer-associated variants we generated, the amino acid change did not affect the overall structure; we did not observe an increase in the insoluble fraction during nucleotide incorporation, and protein production yields were similar to WT, and the secondary structure was unperturbed (Figure 2). Pre-steady state kinetics was our preliminary functional test, and the observed rate of polymerization (*k_obs_*) of the T2161I variant was twice as fast as WT, but still fit to a biphasic equation as seen by many DNA polymerases (Figure 3, Table 2). Our results show that the observed rate under pre-steady state conditions (excess substrate) matches with the polymerization rate (142 ± 22 s^-1^ and 167 ± 12 s^-1^, respectively), but there is a dramatic reduction in affinity for correct nucleotide when paired with templating base G with a K_d(dNTP)_ of 199 μM compared to WT at 5 μM. T2161I prefers to incorporate dT opposite G with a 36x higher affinity. Likewise, T2161I is 38 times less efficient at incorporating correct nucleotide and 77 times less faithful (Table 3).

### E2406K Fingers Domain mutation

The stage IV ocular melanoma mutation located in the fingers subdomain presented similar kinetic behavior to the other cancer-associated variants (Figure 1, Table 1). Secondary protein structure characteristics were unchanged (Figure 2), and under pre-steady state conditions, it too experienced biphasic burst kinetics but at about half the observed rate of WT (Figure 3, Table 2). Interestingly, E2406K polymerization rate was observed to be around 89 s^-1^, this variant also had a reduced affinity for correct nucleotide compared to WT with a K_d(dNTP)_ value of 93μM. Its overall efficiency was also 34 times less than that of WT and had a reduced fidelity of 40 times compared to WT (Table 3). More importantly, this is the only variant that came from a tumor that also had BRCA1 and BRCA2 mutations^54^ suggesting that repair capabilities could further challenged due to a mutagenic and slower Pol θ with the potential to promote drug resistance and tumor growth^55^.

### L2538R Palm Domain mutation

The palm domain variant L2538R came from a melanoma patient with stage IV sun-exposed melanoma (Figure 1, Table 1). Similar to E2406K, this variant had a reduced observed polymerization rate of 27 ± 5 s^-1^ which is half of what was observed with WT. Under single turnover conditions, it remained the slowest of all the variants with a polymerization rate of 41 s^-1^ Despite this, the affinity for nucleotides was more similar to WT, with only a slight reduction in overall K_d(dNTP)_ and was the most efficient of the variants with only a 12-fold reduction in efficiency compared to WT. L2538R had the greatest affinity for dATP with a 34-fold loss in fidelity compared to WT.

In summary, we have biochemically characterized three melanoma-derived mutations in human DNA Polymerase theta. We have demonstrated that these variants experience altered nucleotide incorporation, with a preference for incorrect nucleotide selection compared to WT as observed with the reduced correct nucleotide affinities. Since these variants were derived from cancer patients, we provide evidence for critical residues important for overall DNA repair and genomic stability. Taken together, it could be hypothesized that the tumor utilizes the mutagenic repair pathway of mutated Pol θ to promote genomic instability and further evade cancer therapeutics. Our work identifies novel biomarkers that lay the groundwork for potential treatments of melanoma and factors of tumorigenesis.

## Experimental Procedures

### Materials

All chemicals and reagents were purchased through Sigma-Aldrich (St. Louis, MO), Bio-Rad Laboratories (Hercules, CA) and AmericanBio (Canton, MA) unless indicated. Oligonucleotides were purchased from Integrated DNA Technologies (Newark, NJ). Oligos for assays are purified through HPLC with standard desalting. Activities of wild-type (WT) and variants were assayed at a minimum of three replicates, using at minimum 3 protein preparations, and by at least two individuals.

### Variant acquisition and structural predictions

Melanoma tumor samples were obtained through a collaboration with the Specimen Resource Core of the Yale SPORE in Skin Cancer. Sample preparation, nucleic acid extraction, and whole-exome sequencing were collected as previously described ^54,56^. To assess the impact of each amino acid substitution on overall structure and function we analyzed the full amino acid sequence under default conditions using the algorithms SIFT^41^ and PolyPhen-2^42^. All five variants were predicted to alter protein structure and thus function and were selected to be biochemically analyzed.

### Generation of cancer-associated variants

pSUMO3 vector containing truncated c-terminal wild-type human DNA Polymerase θ (POLQM1^28^) was generously donated by Dr. Sylvie Doublié from the University of Vermont.

Mutations were generated by site-directed mutagenesis (table 2) on this plasmid using QuikChange II Site-Directed Mutagenesis Kit (Agilent Technologies) and plasmids were verified through sequencing through the Molecular Informatics Core at the University of Rhode Island.

**Table 2.**
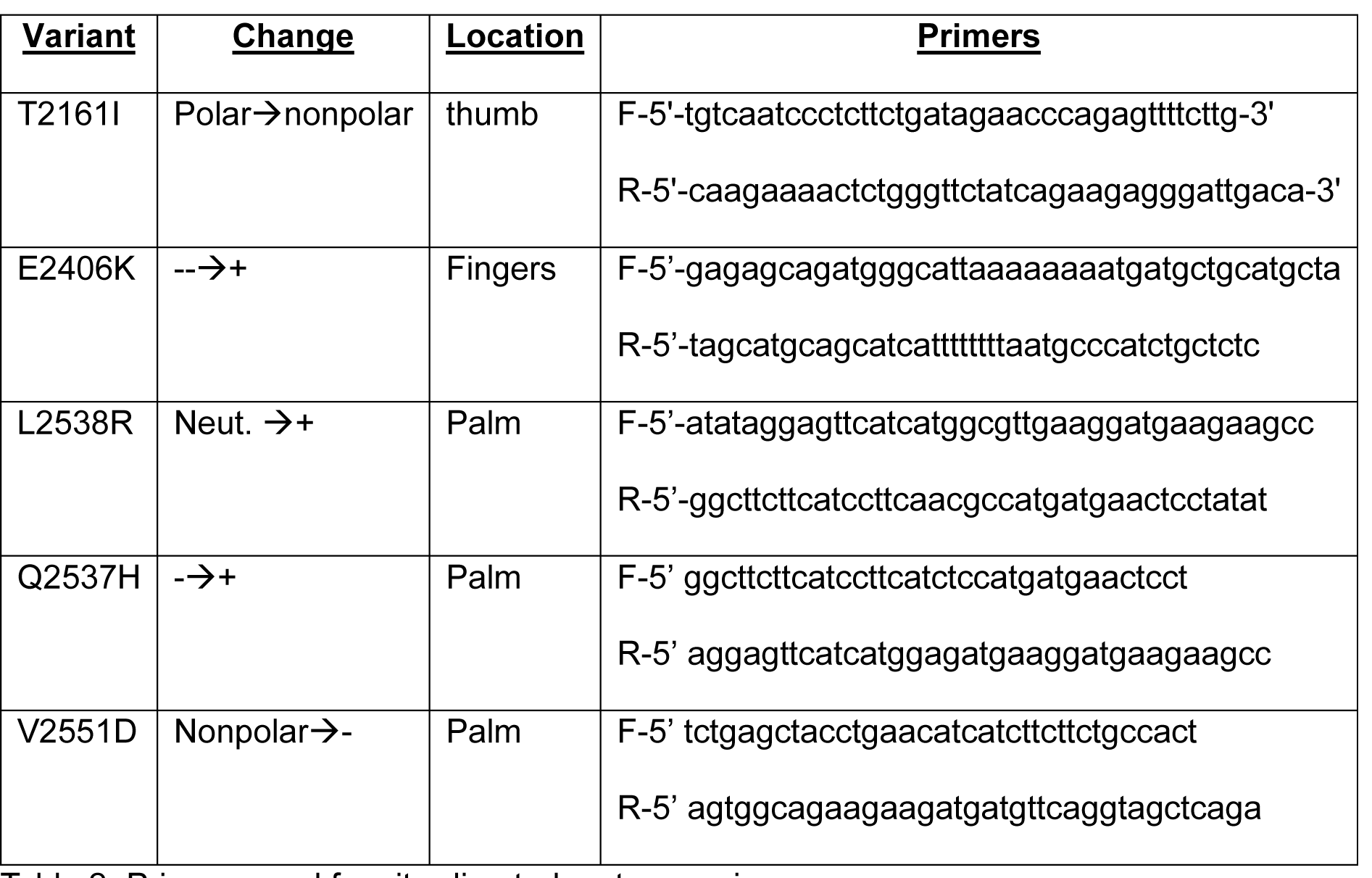
Primers used for site-directed mutagenesis.

### Molecular Modeling

PyMol 1.3^57^ were used to generate a representation of the c-terminal POLQM1 and its cancer-associate variants from the crystal structure as previously presented^26^.

### Expression and Purification of WT Pol θ and its Cancer-Associated Variants

Expression and purification of plasmids containing human WT (WT) and/or its variants followed a modified protocol similar to previously described^28^ but optimized for maximum DNA polymerase activity. Plasmid was transformed into Rosetta2(DE3) pLysS competent cells (EMD Millipore) and plated on LB agar plates containing 100μg/mL ampicillin and 34μg/mL chloramphenicol. Colonies were directly inoculated into 1L of Terrific Broth and incubated for 60 hours at 20°C. Cells were harvested by centrifugation at 5K RPM for 10 minutes and stored at - 80°C.

Pellets were thawed on ice and resuspended in Lysis Buffer (20mM Tris, pH 7; 300mM NaCl; 0.01% NP-40; 10% glycerol; 20mM Imidazole; 5mM BME; 0.120mM PMSF; and EDTA-free Protease Inhibitor Mixture (Roche Applied Science)) and sonicated for 6-8 rounds for 30 seconds. Soluble cell fractions were separated via centrifugation at 15K RPM for 30 minutes. Soluble protein fractions were separated by fast protein liquid chromatography (FPLC) with an imidazole gradient by mixing binding buffer A (20mM Tris, pH 7; 300mM NaCl; 0.01% NP-40; 10% glycerol; 20mM Imidazole; 5mM BME) with elution buffer B (buffer A with 500mM Imidazole) on a 5mL His-Trap FF Crude Nickel Column (GE Healthcare). Fractions containing Pol θ were pooled and were separated further on a 5mL HiTrap Heparin HP (GE Healthcare) with a NaCl gradient by mixing binding buffer C (20mM Tris, pH 7; 300mM NaCl; 0.01% NP-40; 10% glycerol; 5mM BME) and elution buffer D (buffer C with 2M NaCl). Pooled fractions containing Pol θ were cleaved overnight at 4°C with SUMO2 protease (Fisher Scientific).

Cleaved protein was separated by a 5mL HiTrap Chelating HP (GE Healthcare) column using an imidazole gradient by mixing binding buffer E (20mM Tris, pH 7; 300mM NaCl; 0.01% NP-40; 10% glycerol; 10mM Imidazole; 5mM BME) with elution buffer B. Cleaved Pol θ was collected in the flow-through fraction with the 6x-sumo remaining on the chelating column. The imidazole was removed from the protein preparation by a final HiTrap Heparin column by mixing buffer C and buffer D omitting NP-40 detergent. Cleaved, purified protein (yield approximately 10μM) was rapidly frozen in liquid nitrogen and stored at -80°C for approximately 3 months.

### Generation of DNA substrate

The DNA substrate was generated using two oligodeoxynucleotides (IDT). The primer (5’6-FAM label) was annealed to the complementary 40-mer template as described below:

5’-/FAM/ TTTGCCT TGA CCA TGT AAC AGA GAG

CGGA ACT GGT ACA TTG TCT CTC <u>G</u>CA CTC ACT CTC TTC TCT

Annealing was verified by a 12% native PAGE with annealed and primer only samples and scanned on an RB Typhoon scanner (Cytiva) with the FAM fluorescence filter.

### Circular Dichroism

To compare secondary structure of WT to Pol θ variants, the ellipticity of 3μM of protein 10mM K_2_HPO_4_ were measure from 190 to 280nm at room temperature (20°C). Samples were measured in a 0.2cm quartz cuvette in J-815 CD Spectropolarimeter (Jasco).

### Rapid Chemical Quench Assays

100nM Pol θ was pre-incubated on ice with 300nM 5’FAM labeled DNA substrate and rapidly mixed with 100μM dCTP (correct nucleotide) and 10mM MgCl_2_ in reaction buffer (20mM Tris HCl, pH 8.0, 25mM KCl, 4% glycerol, 1mM BME, and 80μg/mL BSA) using an RQF-3 Chemical Quench Flow apparatus (KinTek) at 37°C from 0-0.6s. Reactions were stopped by the addition of 0.5M EDTA and collected into microcentrifuge tubes containing 90% formamide sequencing dye. Products were separated on 15% denaturing polyacrylamide gel and scanned on an RB Typhoon scanner as described above. The extended products (n+1) were quantified using ImageQuant software and data fitted to a non-linear regression biphasic burst equation using Prism 9 GraphPad Software

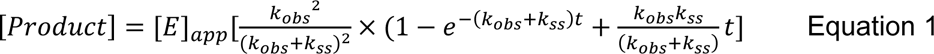

*Active Site Titration* The DNA substrate was titrated from 0 to 300nM against 100nM Pol θ, 100μM dCTP, 10mM MgCl_2_ in reaction buffer at 37°C from 0-0.6s. Reactions were stopped by the addition of EDTA and scanned on the Typhoon as described above. Products were quantified and data fit to Equation 2. The E_app_ was plotted against concentration of DNA. The concentration of Active Sites and K_D(DNA)_ were determined by fitting data to a quadratic equation^58^.

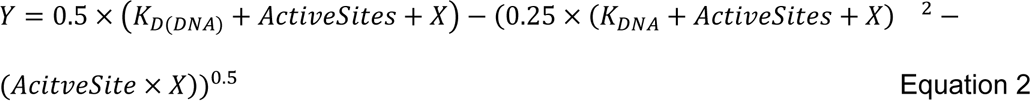

*Empirical Enzyme Titration* To ensure single turnover conditions, a primer extension assay was performed at varying ratios of protein:DNA. DNA substrate (50nM) was reacted with varying concentrations of Pol θ (50nM to 500nM) with 100μM dCTP, 10mM MgCl_2_, in reaction buffer.

Products were separated on a 15% denaturing polyacrylamide gel and scanned on an RB Typhoon as described above. Data was fitted to a single exponential equation

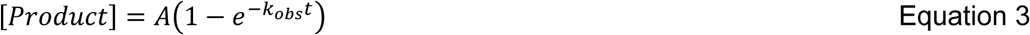

*Single-turnover kinetics* Pol θ and 5’FAM labeled DNA substrate were assayed at a 4:1 ratio as determined by the active site and empirical enzyme titrations. Correct nucleotide, dCTP, was titrated from 0 to 1000μM with 50nM DNA substrate and 200nM Pol θ from 0 to 0.6s on the RQF at 37°C. Incorrect nucleotides were titrated from 0-1000μM from 0 to 300s with the same Pol θ and DNA concentrations by hand at 37°C. Products were separated on a 15% denaturing polyacrylamide gel and scanned and quantified as described above. Data was fitted to a single exponential equation 4 to determine *k_obs_* which is the observed rate at each dNTP concentration. *k_obs_* was plotted against nucleotide concentration for each of the 4 deoxyribonucleotides and fitted to the hyperbolic equation:

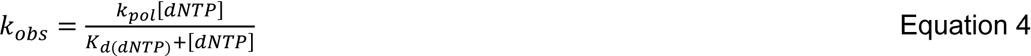

## Acknowledgements

We would like to thank Sylvie Doublié for the generous donation of the plasmid used in these experiments. A special thank you to Ruth Halaban and Antonietta Bacchiocchi for the POLQ melanoma mutations and continued support. Thank you to the various undergraduate chemistry majors at Rhode Island College who participated in the initial set up of this project.

## Funding

Research reported in this publication was supported by the National Institute of General Medical Sciences of the National Institutes of Health under grant number R15GM144903-01 and in part by the Rhode Island Institutional Development Award (IDeA) Network of Biomedical Research Excellence under P20GM103430.

The content is solely the responsibility of the authors and does not necessarily represent the official views of the National Institutes of Health

## Data Availability

All data presented in this manuscript is available upon request to

## Supporting information

N/A

## Conflict of Interest

The authors declare that they have no conflicts of interest with the contents of this article.

## Notes

### Competing Interest Statement

The authors have declared no competing interest.

